# Memory for stimulus duration is not bound to spatial information

**DOI:** 10.1101/2020.07.31.230466

**Authors:** Wouter Kruijne, Christian N. L. Olivers, Hedderik van Rijn

## Abstract

Different theories have been proposed to explain how the human brain derives an accurate sense of time. One specific class of theories, intrinsic clock theories, postulate that temporal information of a stimulus is represented much like other features such as color and location, bound together to form a coherent percept. Here we explored to what extent this holds for temporal information after it has been perceived and is held in working memory for subsequent comparison. We recorded EEG of participants who were asked to time stimuli at lateral positions of the screen followed by comparison stimuli presented in the center. Using well-established markers of working memory maintenance, we investigated whether the usage of temporal information evoked neural signatures that were indicative of the location where the stimuli had been presented, both during maintenance and during comparison. Behavior and neural measures including the contralateral delay activity, lateralized alpha suppression and decoding analyses through time all supported the same conclusion: the representation of location was strongly involved during perception of temporal information, but when temporal information was to be used for comparison it no longer showed a relation to spatial information. These results support a model where the initial perception of a stimulus involves intrinsic computations, but that this information is subsequently translated to a stimulus-independent format to be used to further guide behavior.

## Introduction

Most of our behavior benefits from an accurate sense of time, from holding a fluent conversation with well-timed pauses, to playing music or sports, to navigating traffic. Timing is crucial to determine when to act or when to expect an upcoming event. Therefore, understanding the mechanisms, processes and representations of time and durations in the brain are essential aspects of understanding human cognition and behavior as a whole. To this end, a wide array of competing models and theories have been proposed: One of the most recent review articles lists eighteen different models (Addyman et al., 2016; see also Hass & Durstewitz, 2014), each relying on different assumptions and computations.

Models of timing are typically evaluated and constrained on the basis of errors and biases in time perception displayed by both humans and other animals. For example, the ‘scalar property’ describes that variability in perceived duration is proportional to the objective duration (Gibbon, 1977; Hass & Herrmann, 2012; Kacelnik et al., 1990; Malapani & Fairhurst, 2002). As this is so commonly found in both human and nonhuman timers this has been described as a hallmark of interval timing models (Buhusi & Meck, 2005; Okamoto & Fukai, 2001). Other factors that are typically considered include how time perception is affected by the physical properties of the timed stimulus (Eagleman, 2008; Schlichting, de Jong, et al., 2018; Walsh, 2003), the contextual or emotional salience of the stimulus (Allman et al., 2014; Droit-Volet & Meck, 2007; Ernst et al., 2017; Halbertsma & van Rijn, 2016; Matthews, 2011), the influence of past timing experiences (Jazayeri & Shadlen, 2010; Lejeune & Wearden, 2009; Maaß, Schlichting, et al., 2019; Roach et al., 2017; Schlichting, Damsma, et al., 2018; Taatgen & van Rijn, 2011), and how temporal percepts are affected by neuropharmacological substances (Coull et al., 2011; Meck, 1996; Soares et al., 2016) or aging processes (Lustig & Meck, 2001; Maaß, Riemer, et al., 2019; Turgeon et al., 2016).

While such studies have yielded important constraints on the dynamic computations that underlie a temporal percept, much fewer studies have directly investigated the actual representation of this percept. Consider, for example, the simple task of sequentially perceiving two intervals of different durations, followed by the question: “which interval lasted longer?” Such a task will not only involve the perception of time, but also requires one to commit some representation of the first duration to memory in a manner that it is then usable for comparison with the second interval. The present study aims to investigate the nature of this representation.

With respect to this representation, the wealth of models and theories on timing can be coarsely divided into two classes. In *dedicated* clock models (e.g., Gibbon, 1977; Gu et al., 2015; Matell & Meck, 2004; van Rijn et al., 2014), the representation of time is functionally decoupled from the imperative stimulus. These models assume that sensory events are processed and subsequently fed into a largely independent timing system. Sensory information serves to signal the onset or offset of an interval, and can, in some models, trigger or modulate internal dynamics that give rise to the temporal percept. However, the end result of timing is represented by a dedicated circuit and is in itself not linked to the presented stimulus or any of its features. For example, the Striatal Beat Frequency model assumes that the representation of time derives from a concert of oscillatory activity at slightly different frequencies, which are reset by the perceived onset of an interval. Whenever time is to be read out and committed to memory, it is represented by the state of the peaks and troughs in each frequency band. This representation is completely independent from the stimulus that initially triggered the oscillations.

By contrast, *intrinsic* clock models (e.g., Finnerty et al., 2015; French et al., 2014; Mauk & Buonomano, 2004; Paton & Buonomano, 2018) assume that the representation of time is a product of the stimulus percept itself. In these models, perceiving the stimulus triggers dynamics that are intrinsically part of a stimulus representation, just like features such as location and shape, but that can be used to infer time. In some models, such as the Temporal Context Model (Shankar & Howard, 2010; and its successors Shankar & Howard, 2011, 2013) and the Gaussian Activation Model of Interval Timing (French et al., 2014), the perceived time of an interval is directly derived from its representation in memory. These models typically assume that the stimulus representation is not static but gradually changes as a function of time, which endows these representations with the capabilities of an intrinsic clock.

While dedicated and intrinsic clock models differ in their assumptions regarding the representations and dynamics involved in the perception of duration, models of both classes assume that in a comparison task, the duration percept must somehow be committed to memory. As such, the nature of working memory for time poses important implementation constraints for all models of timing. Nevertheless, models typically make no claims about how temporal information is represented in working memory. Therefore, here we attempt to map this representation by means of electroencephalography (EEG). In particular, we will measure lateralized signals that are ubiquitously studied in visual working memory research and which suggest that space might be a crucial dimension to bind features of representations in memory. Here, we investigate whether these spatial signatures can also be observed when durations are to be maintained.

The first of these signatures is the Contralateral Delay Activity (CDA), a component in the event-related potential (ERP) that was originally identified in change detection experiments (Eimer & Kiss, 2010; Ikkai et al., 2010; Luria et al., 2016; Vogel & Machizawa, 2004). When participants are required to remember items presented on one side of the screen, then a sustained occipital contralateral negativity is observed. The amplitude of this component can be related to the memory load on the trial, and to the memory capacity of the participant. Interestingly, a CDA is also found in studies where the location of the remembered stimulus is irrelevant for the upcoming task, for example when the stimulus is to be used in visual search (Carlisle et al., 2011; Woodman et al., 2013) or merely has to be recognized in the center of the screen (Gunseli et al., 2014). Together with the CDA-component found in the ERP, working memory for laterally presented items is often found to yield a contralateral suppression of frequency power in the alpha band (Klimesch, 2012; Mazaheri, 2010; Van Driel et al., 2017). Such lateralized alpha suppression has been linked to both maintenance of items in working memory, as well as covertly attending a location in space (Foster et al., 2017; Sauseng et al., 2005; van Diepen et al., 2016; van Moorselaar et al., 2018).

Lateralized neural signatures have not only been found for memory *maintenance* but similarly for when visual information is *retrieved* from memory. The N2pc is an occipital contralateral negative inflection of the ERP typically found 200–350ms after a stimulus and is assumed to reflect attentional orienting (Eimer, 1993; Eimer & Grubert, 2014; Luck et al., 1993; Luck & Hillyard, 1994; Tan & Wyble, 2015). While the N2pc is classically studied in the context of visual search, various studies have reported that retrieving lateralized stimuli from working memory similarly evokes an N2pc (Dell’Acqua et al., 2010; Kuo et al., 2009; Leszczyński et al., 2011). Much like these effects on the ERP, orienting to endogenous representations in working memory has been found to produce lateralized suppression of alpha-band power. In these cases, alpha power is found to be suppressed, contralateral to the side that an item that was presented that either has just become relevant or is expected to become relevant. In most studies such orienting is explicitly triggered by means of retro-cues (Poch et al., 2014; van Ede, 2018; Wolff et al., 2017), but it is also observed in experiments where a sequence of subtasks sequentially requires the activation of a different item from memory (de Vries et al., 2017, 2019). Particularly relevant to the present research, orienting responses in the alpha band have also been found in a task where no explicit cues for retrieval are given, but where instead the duration of the memory delay itself informed participants which of two memory items is more likely to be tested (van Ede et al., 2017).

These findings constitute lateralized signatures of maintenance and retrieval of working memory items. Crucially, in most of these studies, lateralized EEG-responses were found despite the fact that spatial information was itself irrelevant for the task: often only non-spatial features of these items, such as color or orientation needed to be maintained. These findings suggest that the memory representation of visual features inherently includes spatial information, which is subsequently detected in the EEG. The visual cortex is spatiotopically organized, and space is the primary dimension along which features are bound in many computational models of working memory storage (Oberauer & Lin, 2017; Schneegans & Bays, 2017; Swan & Wyble, 2014). Therefore, we reason that if the working memory representation of time is similarly bound to sensory information, these spatial neural signatures would be the most likely markers to detect such binding.

Specifically, participants were sequentially presented with two intervals (Interval 1 and Interval 2), separated by a memory delay. Participants subsequently had to determine which of the intervals was longer. Each interval was presented by means of visual markers: stimuli that were briefly presented to indicate the start- and the end-signal for timing. Critically, for Interval 1 these markers were lateralized, that is, presented on either the left or right side of the display, whereas the markers for Interval 2 were both presented in the center. If temporal information uses a representation that is bound to spatial information, then we should find lateralized EEG-signals indicative of maintenance, retrieval, or anticipation during the centrally presented second interval. In that case, their dynamics and their timing could be informative of how such a comparison task is solved: for example, temporal information might be retrieved either only at the start or end of Interval 2, or might be held active throughout comparison. Furthermore, signals of retrieval might reflect *anticipation* of the start- or end of Interval 2 (cf. van Ede et al., 2017), or they might be evoked *in response* to the stimuli marking its duration (cf. Kuo et al., 2009).

To further disentangle the precise role of memory representations in such a comparison task, we additionally manipulated the manner in which Interval 1 was presented. In blocks with ‘Same’ trials, the start- and end-marker were presented in the same side of the screen, whereas in ‘Opposite’ blocks, they were on opposite sides. If we observe lateralized neural responses in both of these block types, then the direction of such lateralization could help us identify whether this reflects retrieval of the start-moment, the end-moment or both. Should we only observe lateralization in the EEG in ‘Same’ blocks, then this might point to a representation where the interval duration as a whole is bound to one location in space, and retrieved during comparison.

Additionally, we explored whether lateralized neural responses would differ between correct- and incorrect trials, thereby attempting to relate these signatures to behavior. If the representations underlying lateralized neural signatures play a functional role in maintaining accurate memory for time, then these neural responses may be found to be weaker or absent on incorrect trials. However, as we will show, behavioral performance on this task was reasonably high. As a result, the number of trials to assess neural signatures on ‘incorrect’ trials was limited and variable across participants.

## Method

### Participants

We collected data from 24 healthy participants with normal or corrected-to-normal vision, who were recruited through the participant pool of the Faculty of Behavioral and Movements Sciences of the Vrije Universiteit Amsterdam. All participated for course credits or monetary compensation (€10/hour). Data of four participants were discarded after EEG data preprocessing (described below), two of which due to the number of noisy channels (8 and 10 electrodes) and two due to the high percentage of discarded data segments (38% and 52%) due to contamination from muscle artefacts or horizontal eye movements. No participants were excluded based on behavior. We inspected behavioral performance, we fit an individual logistic regression model to the proportion of ‘longer’ responses as a function of Δ*t* = (Interval 2 - Interval 1), and all slope coefficients were less than 2 SD away from the group mean. The final sample contained 20 participants (ages 19–26, Mean age 21.9, 10 female). For all participants, informed consent was obtained before participation, and all procedures during the experiment were in accordance with the Helsinki declaration. The protocol was approved by the ethical review board of the Faculty of Behavioral and Movement sciences of the Vrije Universiteit Amsterdam.

### Procedure and stimulus presentation

Participants were seated in a darkened, sound-attenuated room at 75cm viewing distance from a 22 inch screen (Samsung Syncmaster 2233, 1680 × 1050 resolution, 120 Hz refresh rate). The experiment was programmed and presented using OpenSesame (Mathôt et al., 2012) with the PsychoPy back-end (Peirce, 2007). The stimulus sequence of a trial is schematically depicted in Figure 1A (gray and black colors in Figure 1A are inverted for visibility). Trials started with a gray fixation cross (0.2°), on a black background for 1000ms, followed by the onset of three, horizontally aligned gray placeholder circles (radius 1.97°, one in the center and two at 9.83° eccentricity) around a central gray fixation dot (0.2°). The placeholders and fixation dot stayed on screen until the end of the trial.

**Figure 1.**
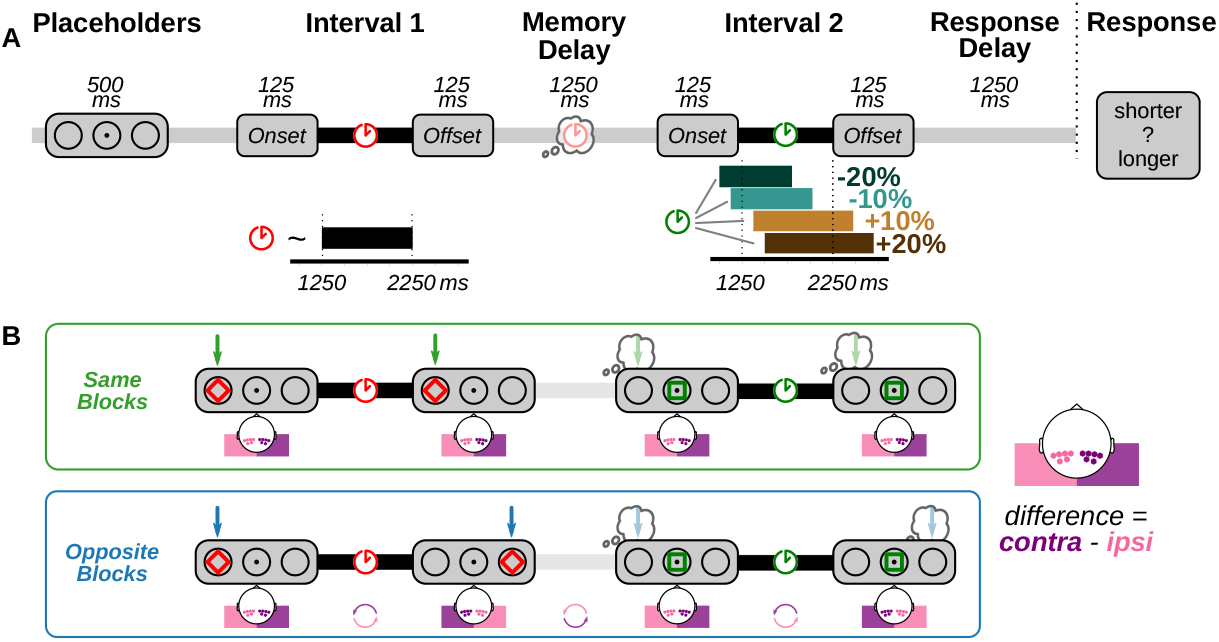
Schematic representation of the design. Note that in the experiment, fixation-, placeholder stimuli were gray on a black background (here inverted for legibility) **A** The general sequence in all trials. A trial is initiated by the appearance of three placeholder stimuli, indicating where Onset- and Offset markers can appear. This is followed by the presentation of Interval 1, the memory delay, Interval 2 and the response delay, after which participants are probed to indicate which Interval was longer. The duration of Interval 1 (red clock) was a uniform random sample between 1250 and 2250. Interval 2 duration (green clock) was based on Interval 1, and was either 10% or 20% shorter or longer. **B** Two trial types were presented in blocked fashion and differed in how Interval 1 was presented. Both markers in ‘Same’ blocks (top row) were in the same location, but in ‘Opposite’ blocks (bottom row) they were in opposite locations. Note that the marker locations in Interval 1 determine which electrodes are labeled as ‘contralateral’ and ‘ipsilateral’ in the analyses, both for Interval 1 as for Interval 2 where markers were presented in the center. For ‘Opposite’ trials, this means that these labels alternate for each consecutive data epoch in the same trial, as illustrated in the bottom row.

The presentation of Interval 1, the ‘standard interval’, started 500ms after placeholder onset, and was indicated by an onset- and offset marker flashing (125ms) in either the left or right placeholder. Both markers were red diamonds (3.94° width and height). Their SOA defined the standard interval, which was randomly sampled from a uniform distribution 𝒰 (1250 *-* 2250*ms*). After the offset marker was presented, a 1250ms memory delay followed. Interval 2, the ‘comparison interval’, was then similarly presented by means of an onset- and offset marker, each a green square (same surface area as the diamonds, sides 2.79°; 125ms) presented in the central placeholder circle. The duration of Interval 2 was derived from the sampled duration of Interval 1, factorially defined to last either 10% or 20% shorter or longer. For example, if the sampled duration of Interval 1 was 1500ms, Interval 2 could last either 1200, 1350, 1650 or 1800ms. The resulting uniform distributions of possible Interval 2 durations are depicted on the right of Figure 1A. Visualizations of the empirical interval distributions are available in the online Supplemental Information (https://osf.io/7gpka/, Figure S1).

The offset marker of Interval 2 was followed by a 1250ms response delay. The shape and color of the interval markers were chosen so that Interval 1 and Interval 2 would not share any non-temporal features that might automatically trigger the retrieval of spatial information. The minimum duration between stimuli, 1250ms, was chosen to be long enough for the relatively sluggish lateralized alpha response to emerge and resolve completely (de Vries et al., 2018; Van Driel et al., 2017).

Participants were instructed to maintain fixation on the center of the screen throughout the trial and not to move their eyes towards the peripheral stimuli. They were asked to compare the duration of the stimuli and determine whether the second interval was shorter or longer than the first. Once the response screen appeared, they could indicate their answer by pressing ‘Z’ or ‘M’ on a standard QWERTY keyboard with their left or right index finger. On each trial, the mapping of these keys varied unpredictably and was unknown to the participant until the response screen appeared. This was done to prevent any (lateralized) motor preparation signals contaminating the EEG. On the response screen, the options (Z and M) were presented centrally above or below fixation accompanied by the words “shorter” (above fixation) or “longer” (below). Participants were instructed to prioritize accuracy over response speed.

Participants completed eight blocks of 40 trials each. Before each block, participants were informed how Interval 1 would be presented: via markers on the ‘Same’ side or on ‘Opposite’ sides (Figure 1B). Note that in either block type, participants could not predict the location of the Interval 1 onset marker, but after its presentation, the location of the offset marker was fully predictable. All combinations of the percentage change and the onset location of Interval 1 were presented five times per block in a random order. Block types (Same/Opposite) alternated and their order was counterbalanced across participants. Before starting the experiment, participants completed practice blocks with Same and Opposite marker presentations, ten trials each. These practice trials were not considered in any of the analyses.

### EEG acquisition and data cleaning

EEG data were recorded at 512Hz from 64 channels (BioSemi, Amsterdam, The Netherlands; ActiveTwo system, 10–20 placement; biosemi.com), with two additional electrodes placed at the earlobes, two placed 2cm above and below the right eye, and two electrodes placed 1cm lateral to the external canthi. All offline analyses and data cleaning steps were performed using MNE-python (Gramfort et al., 2013; Gramfort et al., 2014) and R (R Core Team, 2018). EEG data were re-referenced to the average of the data from the earlobes, and V/HEOG traces were created by subtracting data from the opposing channels around the eyes.

For data cleaning, three filtered versions of the raw dataset were created by means of zero-phase FIR filters: band-passed at at 110–140Hz to highlight muscle artefacts; high-passed at 1Hz, removing medium-slow drifts to be used for independent component analysis (ICA); and one high-pass filtered at 0.1Hz to be used for the main analyses. Most algorithms that were used for data cleaning assume epochs of equal length reflecting “trials” as their input. Unless otherwise specified, we used “preprocessing epochs” as input for these algorithms. The size of these epochs was based on the shortest possible interval durations (1250ms and 1000ms for Interval 1 and 2 respectively), the fixed 1250ms memory delay between them, and the fixed 1250ms response delay. These epochs thus spanned −2500ms – 2250ms around the onset of Interval 2. The preprocessing epochs were not used in any of the analyses, which instead were based on four epochs surrounding the on- and offset markers (see below). Rather, preprocessing epochs were used to determine the thresholds for artefact detection, which were subsequently used to accept or reject data in the analysis epochs.

Following the PREP-pipeline (Bigdely-Shamlo et al., 2015) we first identified excessively noisy channels by means of the RANSAC algorithm, as implemented by the ‘autoreject’ package (Jas et al., 2017). This procedure generates permutations of the epoched data, and predicts full channel activity by interpolating data from a subset (25%) of the channels. If the correlation between the observed and interpolated data is less than a threshold value (r < 0.75) in more than 40% of the epochs, the channel is classified as ‘bad’. In our dataset, the algorithm identified 6 (n=1), 4 (n=1), 2 (n=2), and 1 (n=3) faulty channels per participant, which were found to be in agreement with visual identification. Faulty channels were primarily located on more peripheral electrode sites, and did not overlap with the preselected electrodes of interest at parieto-occipital sites (see Supplemental Information, Table S1). We excluded these channels from all other preprocessing steps.

Muscular activity can introduce broadband noise that overlaps with neural activity in the EEG signal, which can be challenging to filter out (Muthukumaraswamy, 2013). Therefore, we sought to identify data segments that might have been contaminated by muscle artefacts and remove data from epochs that contained such contaminated segments. To do so, we used a procedure adopted from the PREP-pipeline (Bigdely-Shamlo et al., 2015) also offered by FieldTrip (Oostenveld et al., 2011). This procedure is based on the observation that muscle contamination is characterized by high-frequency power found simultaneously across multiple electrodes. To identify epochs with such contamination, we used the 110– 140Hz band-passed dataset, computed its Hilbert envelope and convolved the result with a 200ms boxcar averaging window. This yielded a per-channel time course estimate of high-frequency power in the original signal. Across all data in the preprocessing epochs, we computed a per-channel median and median absolute deviation, and used these to compute a robust Z-score for all data inside and outside these epochs. Data at time points where the Z-score averaged across channels exceeded 5.0 were marked as contaminated and were not considered in future analyses.

ICA was used to identify and remove artefacts in the data caused by eye blinks. We first subsampled the high-pass filtered (at 1Hz) dataset at 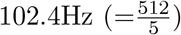, and computed independent components (using extended-infomax ICA, the default in EEGLAB; Delorme & Makeig, 2004). From the resulting spatial components, we visually identified those corresponding to blinks. To confirm the validity of this approach, we defined ‘blink epochs’, as 1000ms windows around local maxima in the 1–10Hz band-passed VEOG signal. The time course of the selected ICA components, and only these components, had a high correlation with the VEOG signal in these epochs (*r*^2^ > .5).

To identify horizontal eye movements, we used a procedure inspired by methods from ERPLAB (Lopez-Calderon & Luck, 2014): Data from the HEOG signal were high-pass filtered at 1Hz, and convolved with a stepwise kernel, defined by 150ms each of −1 and +1 values, with a 25ms linear ramp between them. In the resulting signal, local maxima that exceeded the 99th percentile were marked as potential horizontal saccades. Data from the start of such a mark up to the end of the current trial were excluded from further analyses.

The autoreject algorithm (Jas et al., 2017) was used on the preprocessing epochs in order to identify trials with unreasonably high amplitude fluctuations. This algorithm improves on typical epoch rejection methods that use a fixed threshold, and instead estimates optimal channel-specific thresholds by means of cross-validation methods inspired by RANSAC. In epochs where a channel’s peak-to-peak value exceeds its threshold, autoreject will initially attempt to interpolate that channel’s data from neighboring channels. Only if the number of to-be-corrected channels in an epoch is above an additionally fit integer *k*, the entire epoch is rejected. We fit autoreject on the preprocessing epochs to find individual thresholds per participant per channel (Supplemental Information, Figure S2)).

Using the results of these preprocessing algorithms, we created ‘cleaned’ data epochs around moments of interest. In each trial, four epochs were defined with data surrounding the presentation of each marker: the onset/offset of Interval 1 and the onset/offset of Interval 2 (Table 1). Time windows were chosen to be maximally long without overlapping with other markers (Table 1). As a consequence, onset- and offset-locked intervals contain partially overlapping data, with the amount of overlap dependent on the interval duration. For each of these time windows, ‘cleaned’ epochs were created: first, the raw dataset (high-pass filtered at 0.1Hz) was loaded after which the ICA components related to blinks were removed. Data epochs were extracted, where epochs were dropped if they contained muscle artefacts or if they contained data following a horizontal eye movement on the same trial. The autoreject algorithm was then applied to interpolate data with extreme peak-to-peak amplitudes or drop rejected epochs. Finally, data from faulty channels as identified by RANSAC were interpolated on the basis of the neighboring channels. The resulting number of clean epochs per participant used in all analyses is given in online Supplemental Information, Figure S3.

**Table 1.** Epoch definition

| Epoch / Event | $t_{\min}$ | $t_{\max}$ | Includes | | |
| --- | --- | --- | --- | --- | --- |
| Onset Interval 1 | -1250 | 1250 | Placeholder onset, | Onset marker, | Start Interval 1 |
| Offset Interval 1 | -1250 | 1375 | End of Interval 1, | Offset marker, | Memory delay |
| Onset Interval 2 | -1375 | 1000 | Memory delay, | Onset marker, | Start Interval 2 |
| Offset Interval 2 | -1000 | 1250 | End of Interval 2, | Offset marker, | Response delay |

### Event related potentials

To construct ERPs, all epochs were baseline corrected by subtracting the average data of each channel in the 100ms leading up to the onset of the placeholders ^1^. We then computed two average signals for the electrode clusters [P7, P5, PO7, PO3, P3, P1] and [P8, P6, PO8, PO4, P4, P2] on left and right parieto-occipital sites. These electrodes were selected on the basis of earlier work (de Vries et al., 2017, 2019; Van Driel et al., 2017), and corresponded to the visually identified locus of peak lateralized responses. On each trial, in each data epoch, these two signals were labeled as ‘contralateral’ and ‘ipsilateral’ with respect to the onset- or offset marker under consideration (Figure 1B). For epochs around the onset- and offset of Interval 2, with centrally presented markers, ‘contralateral’ and ‘ipsilateral’ were defined based on the onset- and offset location of Interval 1. The dependent measure in all ERP analyses was the difference between contralateral and ipsilateral sites, which would indicate lateralized neural responses.

Note that this lateralization definition is in line with convention, but has consequences for how lateralization is computed on ‘Same’ and ‘Opposite’ trials (Figure 1B). To illustrate this point: a ‘Same’ trial where the onset- and offset marker are both presented on the left side, the contrast between contralateral and ipsilateral sites entails computing the difference in activity (Right - Left) in the same way in all four data epochs. However, on an ‘Opposite’ trial, an onset marker on the left is followed by an offset marker on the right, and lateralization in that epoch is computed accordingly (Left - Right). The subsequent data epoch around the central onset marker for Interval 2 is again referenced with respect to the marker location of Interval 1 onset (Right-Left), and data surrounding the offset marker is again referenced in the opposite direction (Left-Right).

### Time-frequency analyses

Time-frequency spectra were computed for the same four data epochs in a trial (Table 1). We computed frequency power for frequencies *F* = 2 to 40*Hz*, in 25 steps on a geometric scale. That is, each subsequent frequency was a multiple of a constant growth factor (1.133Hz). To prevent temporal edge-artifacts, the spectra were computed over epochs that were 1.0s wider than the data of interest. Before computing frequency power, the overall evoked response was subtracted from the individual data epochs. We used multitaper filtering, using 500ms windows tapered with three Slepian windows and subsequently zero-meaned. Data and filters were convolved by multiplication in frequency space after fast-fourier transformation. Note that compared to many approaches using wavelet convolution, this multitaper approach tends to be more accurate in determining the timing of power effects at the cost of being somewhat less accurate in terms of frequency. After filtering, time-frequency data were cropped to align with ERPs, and were sub-sampled to 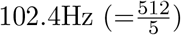. Power was baselined and converted to decibel (dB) with respect to the per-trial average power 300-100ms before placeholder onset.

As for the ERPs, frequency power was spatially averaged across left-and right occipitoparietal electrodes, and subtracting contra- from ipsilateral sites yielded a measure for power lateralization. For analyses contrasting different trial types (same/opposite; correct/incorrect) we focused on the time course of lateralized alpha power, by first averaging data across frequencies between 7 and 13 Hz.

### Classification analyses

Using a classification analysis, we investigated whether information regarding marker locations was maintained or retrieved in a manner that was not reflected in lateralized parieto-occipital ERPs. To this end, we trained logistic classifiers to predict the location of the onset- or offset marker from the 64-channel EEG data. As input to these classifiers, we used the epoched data, low-pass filtered at 35Hz and subsampled to 64Hz. Classifiers were trained on 125ms sliding window segments of data (8 time points × 64 channels) which were Z-scored and used as input features. This setup was constructed to produce high classification performance in decoding the onset marker of Interval 1, and was subsequently applied to decode other epochs. To prevent overfitting, classifier performance in each epoch was evaluated on the basis of 10-fold stratified cross-validation. Performance on each datafold was scored using the area under the curve (AUC) of the receiver-operator curves. Compared to ‘accuracy’, the AUC is a better performance metric to quantify classifier sensitivity while accounting for potential biases in choosing one class over the other. A perfect classification would yield a score of 1.0, whereas scores around 0.5 reflect an inability to meaningfully decode the marker location from the neural data.

### Statistics

Participants’ behavior was characterized by means of General Linear Mixed-Effects regression (Baayen et al., 2008). Models were constructed to predict which of the two intervals was perceived as longer using logistic regression with predictors including: the length of Interval 1, the length of Interval 2, their absolute difference (in seconds), their proportional difference (percentage change), and block type (Same/Opposite presentation). All models with different combinations of these predictors, with and without interactions between main effects, were compared by means of their BIC scores. We report the resulting best model, and report statistical evidence for or against effects (likelihood ratio test *χ*^2^ statistic, corresponding *p*-value, and Δ*BIC*) by comparing nested models that either included or excluded the effect under consideration.

The computation of ERPs, time-frequency- and classification scores were initially all done independently per subject, resulting in time courses and time-frequency spectra for each data epoch under consideration. These multivariate measures were subjected to group-level statistical testing using cluster-based permutation tests (Maris & Oostenveld, 2007). Clusters were defined as regions adjacent in time and frequency where univariate statistical testing yielded a p-value lower than 0.05. Low-variance t-values were corrected for using ‘hat’ variance adjustment (Ridgway et al., 2012) with a correction factor *δ* = 0.001. The t-values within each cluster were aggregated into a single cluster statistic by summing them together. The same cluster statistic was computed in 5,000 random permutations, the results of which were used as a nonparametric null distribution. An observed cluster was considered statistically significant with respect to this distribution at *α* = 0.05.

## Results

### Behavior

The best model in terms of BIC predicted ‘longer’ responses as a function of three additive main effects (Figure 2). First, responses were determined by the percentage change, included in the model as a linear predictor (*χ*^2^(1) = 23.8, *p <* 0.001, Δ*BIC* = 15.0). This predictor captures that participants accurately performed the task: that is, they were more likely to produce a ‘longer’ response on trials that indeed had a longer Interval 2 and vice versa, with more certainty for the 20% change than the 10% change conditions.

**Figure 2.**
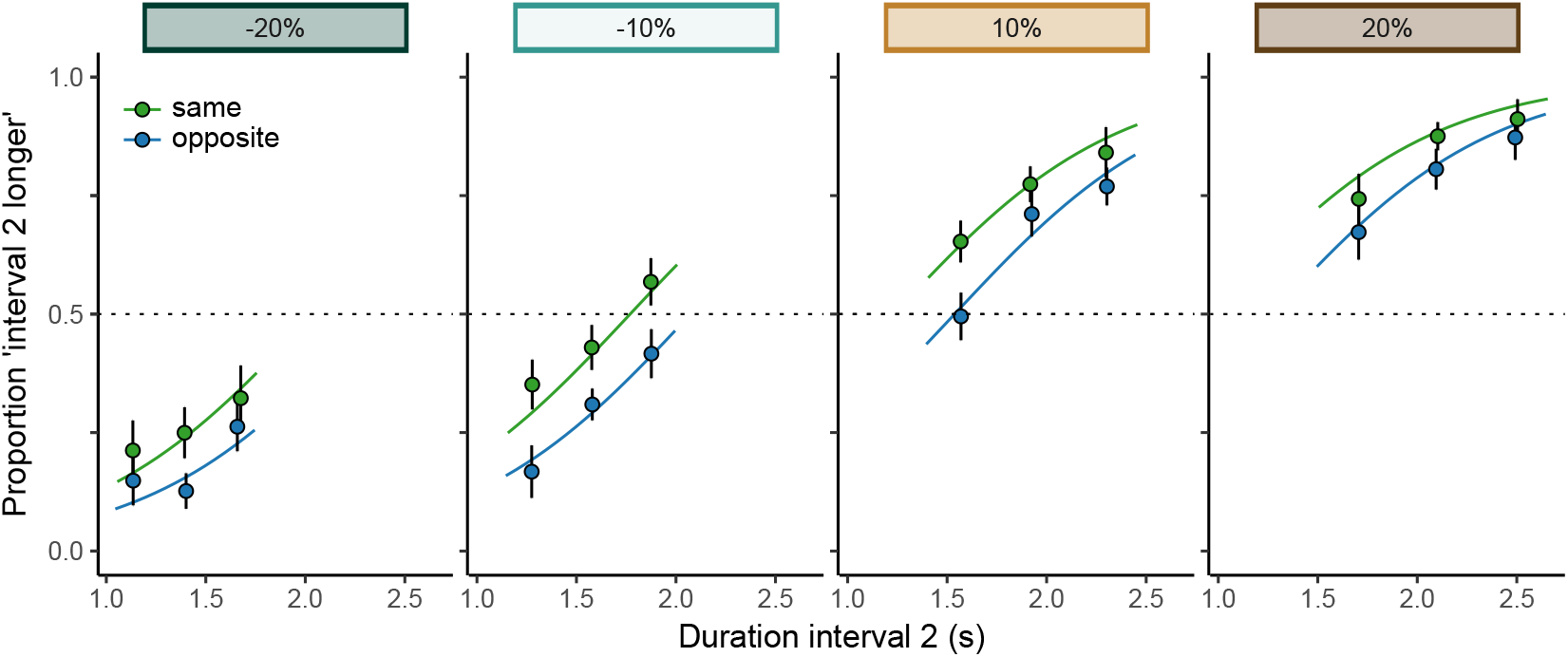
Proportion of trials where participants perceived Interval 2, the comparison interval, as longer. Curves reflect responses predicted by the preferred statistical model with three predictors: the percentage change, Interval 2 duration, and block type (Same/Opposite). For comparison, the data points reflect the proportion of ‘longer’ responses in three ‘Interval 2 duration’ bins defined separately per ‘percentage change’ condition. Error bars reflect 95% within-subject confidence intervals (Cousineau, 2005; Morey, 2008).

Second, participants were more likely to respond ‘longer’ on trials where the second interval was physically long. (*χ*^2^(1) = 298.8, *p <* 0.001, Δ*BIC* = 296.0). This suggests that participants at least in part made a decision based on the absolute duration of Interval 2, regardless of how it related to Interval 1. Although this could reflect participants’ poor memory for Interval 1 on some trials, note that it could also reflect a strategic weighting of evidence: in the present task, the absolute duration of Interval 2 is a good heuristic to determine whether the interval is relatively longer too (see Figure 1A).

Third, we found that participants’ judgments were modulated by the manner of presentation of Interval 1. That is, in blocks where markers were on opposite sides of the screen, Interval 1 was more often perceived longer than Interval 2 than in ‘Same’-blocks (*χ*^2^(1) = 91.2, *p <* 0.001, Δ*BIC* = 82.5). The EEG results presented below offer tentative evidence that this finding relates to the neural response evoked by the offset marker in ‘Opposite’ blocks. However, as this does not immediately relate to the current research question, we have presented a more in-depth investigation elsewhere (Kruijne et al., 2020). In the present article, EEG analyses will focus on neural markers for memory maintenance and retrieval.

### ERP analyses

Figure 3 depicts the ERP waveforms during the epochs surrounding the onset- and offset markers of Interval 1 (A–C) and Interval 2 (D–F). Note that for the centrally presented Interval 2, ‘ipsilateral’ and ‘contralateral’ electrodes are defined with respect to the corresponding marker location during Interval 1 (cf. Figure 1)B.

**Figure 3.**
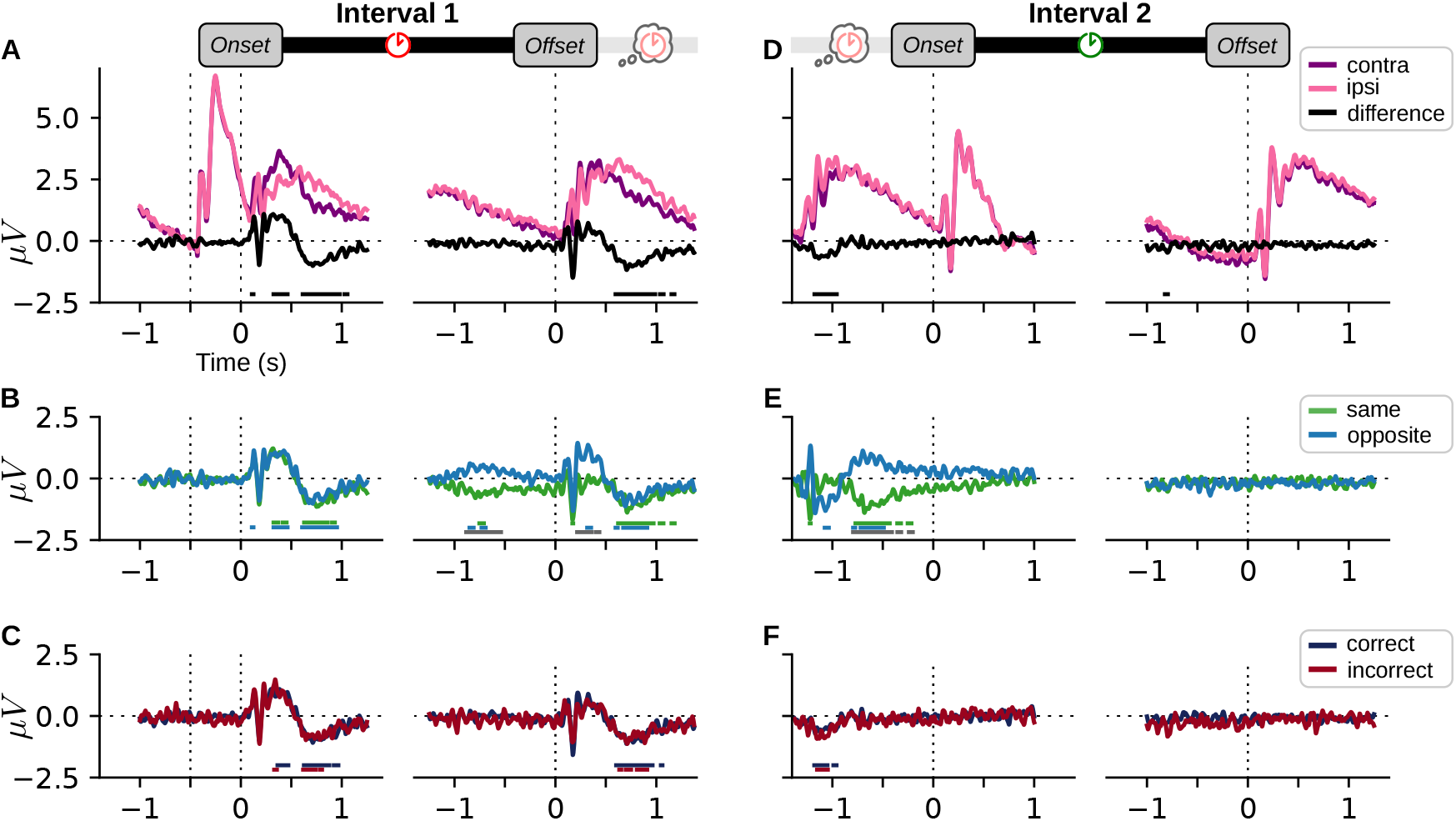
Lateralized ERPs during epochs around the markers of Interval 1 **A-C** and the markers of Interval 2 **D-F**. The three rows of subplots depict, from top to bottom, the ERPs across all trials, in ‘Same’ and ‘Opposite’ trials separately, and in correct versus incorrect trials separately. Data in these plots were low-pass filtered at 35Hz for visualization only. Colored horizontal line segments indicate clusters where the corresponding condition significantly deviates from 0. Gray segments in **B** and **E** indicate significant differences between ‘Same’ and ‘Opposite trials’.

### Interval 1

Figure 3A depicts the ERP computed from all trials at contralateral and ipsilateral sites, as well as the lateralized signal computed as their difference. The figure depicts the ERP evoked by the onset (left) and offset (right) of Interval 1. The left plot shows that before the onset of the interval, the onset of the placeholders (dotted line at *t* = - 0.5*s*) gives rise to a large, bilateral visual response in both hemispheres. The lateralized onset marker at *t* = 0*s* gives rise to a second visual response, which gives rise to lateralization in the ERPs. This is characterized by an early lateralized response, offset by an early (90–145ms) significant postivity. Interestingly, this is shortly after followed by another significant contralateral positive cluster (305–482ms). While we had expected to find a negative component here (the N2pc), there are reported conditions where this component reverses, which we will come back to in the Discussion. After this initial positive component, however, this gradually deflects into a negative component starting at approximately 400ms, yielding a sustained negativity marked by two significant clusters (594–1076ms)^2^. This sustained negativity matches the typical profile of the CDA.

Leading up to the offset of Interval 1 (Figure 3A, right), the grand-average ERP does not show signs of lateralization in anticipation of the upcoming marker. Following the presentation of the offset marker, the grand-average ERP showed a lateralized evoked response that was highly similar to that in response to the onset marker, although here none of the early positive and negative inflections were found to be significant. The later, sustained contralateral negativity resembling a CDA was marked by three significant clusters (576 – 1199ms).

In Figure 3B, the lateralized ERPs are depicted separately for ‘Same’ and ‘Opposite’ trials. These analyses were intended to further separate whether start- or end-marker locations were being maintained or retrieved during comparison, or whether the location was only represented in memory when the entire duration was presented at the same location. In response to the Interval 1 onset marker, both conditions evoke virtually identical ERPs, with similar clusters indicating significant differences in either condition as in the grand-average ERP. Between the two conditions, no significant differences were found.

In the data surrounding the offset marker (right plot), two notable differences between Same- and Opposite presentation conditions can be observed. First, results suggested that the CDA evoked by the onset marker persisted throughout most of Interval 1 in the data before the offset marker. In Same-trials, we still found a contralateral negativity, which gave rise to a significant cluster (−773 to −689ms). In Opposite trials, the ERP is mirrored with respect to the data from the onset marker, and as a result the same CDA component now manifests as a sustained positivity (two significant clusters from −871 to −672ms). Note that because of these mirrored lateralized components, the grand average ERP plotted in Figure 3A shows no significant lateralization as it collapsed across these conditions. The mirrored CDA components also result in a significant difference between the two conditions (−904 to −520ms). No significant clusters were found in the final 500ms of the interval, although numerically, lateralization seemed to persist.

Second, in response to the offset marker, an early lateralized positivity was found for ‘Opposite’ trials (significant cluster 295 to 373ms) but not for ‘Same’ presentations, with a significant difference between them from 195 to 455ms. In part, this condition difference might be accounted for by the CDA difference described above, which numerically persisted up to the offset marker. However, in a separate analysis where these epochs were re-baselined we still found a significant difference. This transient difference therefore probably reflects a stronger visual evoked response to the offset marker in Opposite blocks than in Same blocks. This makes this neural response a prime candidate neural correlate to drive the behavioral bias that intervals with opposite markers are perceived to last longer.

Note that despite this early modulation, there were no apparent differences between Same- and Opposite conditions after approximately 500ms into the memory delay: both showed a sustained CDA of similar amplitude with respect to the location of the offset stimulus which persisted throughout the memory delay.

In the plots in Figure 3C, we depict ERP-traces for correct- and incorrect trials, collapsed across ‘Same’ and ‘Opposite’ conditions. Throughout the epochs surrounding both markers of Interval 1, there were no significant differences between these trials: overall, both conditions closely followed the pattern displayed by the grand-average ERP in Figure 3A.

### Interval 2

Figure 3D depicts the grand average ERP for both the onset (left) and the offset (right) of Interval 2. These markers were all presented in the center of the screen, and lateralized ERPs were computed in relation to the location of the onset- and offset markers of Interval 1 (cf. Figure 1B). While both the onset and the offset marker clearly evoke strong visual responses, the average ERP shows little signs of lateralization, neither in anticipation of the markers nor in response to their presentation. Two significant clusters were found, one early in the memory delay (1,197 to −938ms) and one short-lived negative cluster, long before the offset marker of Interval 2 (−844 and −773ms). However, these are likely to both reflect remnants of the lateralized response to the Interval 1 offset marker, rather than signs of reinstatement or reactivation of a memory representation.

This point is illustrated when lateralized ERPs on ‘Same’ and ‘Opposite’ trials are considered separately (Figure 3E). Note that the ERPs in the time leading up to the offset marker reflect the same data as the data at the end of the offset-epoch plotted in Figure 3B, but with the polarity reversed on Opposite trials. They thus depict the same visual response to the offset of Interval 1, and it is clear to see that the difference in early visual response between the two conditions would give rise to the early lateralized negativity found in the grand-average potential. This early visual response is then followed by a sustained CDA, with opposite polarity for Same- and Different conditions (three significant clusters from - 814ms to −184ms). Qualitatively, however, note that this sustained CDA component appears to persist well into Interval 2 up to 500ms into Interval 2. In the data epoch leading up to the offset marker of Interval 2, this sustained positivity in Opposite trials is expressed as a negativity, again aligning itself with the lateralization of ERPs in Same-trials. This can account for the short lived negativity found in the grand-average data. More crucially though, neither the Same-nor Opposite data showed any signs of ERP lateralization in anticipation of or in response to the centrally presented markers.

The ERPs contrasting correct and incorrect trials again revealed no differences between them, and closely followed the pattern found in the grand-average data.

To conclude, these ERP analyses suggest that actively timing an interval marked by laterally presented stimuli is paired with lateral CDA components resembling maintenance of these markers, suggesting that spatial information concerning these stimuli is stored and maintained in working memory. However, the ERPs gave no indication that this memory representation is then subsequently used at retrieval, that is, during the centrally presented comparison interval.

### Time-frequency decomposition

Figure 4A depicts the full spectrum of lateralized power in data surrounding the onset- and offset of Interval 1. Both markers give rise to large, significant lateralized clusters indicating contralateral power suppression. Both clusters were centered around the alpha band, but extended to frequencies from approximately 5 to 20Hz. This wide spread in terms of frequencies is most likely due to the multitaper approach used here, which has a high temporal accuracy, sometimes at the cost of spectral bleed. The alpha suppression in response to Interval 1 onset was triggered almost immediately after stimulus onset, and spanned approximately 750ms. Around the offset of Interval 1, similar contralateral suppression in the alpha band was found. Interestingly, such suppression already arose during the interval in the time leading up to the offset marker, with data from 500ms before the marker included in the significant cluster. These results might therefore reflect attentional shifting to the predictable location of the offset marker in anticipation of its presentation.

**Figure 4.**
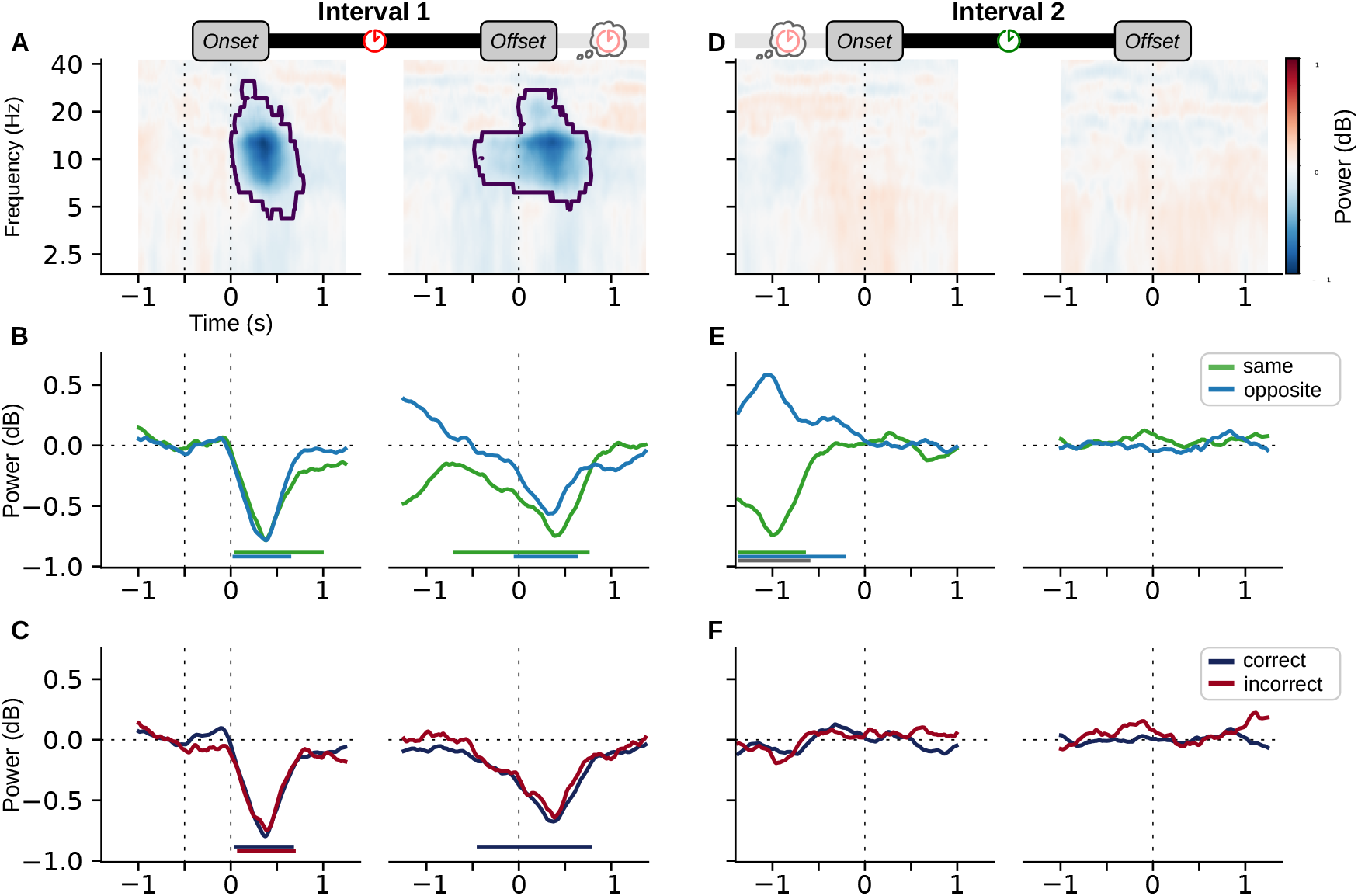
Lateralized Time-frequency decomposition for the four consecutive epochs, plotted as in Figure 3: **A-C** reflect data surrounding the markers of Interval 1, **D-F** reflect data surrounding Interval 2. The three rows of subplots depict, from top to bottom, the Time-frequency decomposition across all trials, alpha lateralization in Same and Opposite trials separately, and alpha lateralization in correct versus incorrect trials separately. Clusters indicating significant lateralization are marked by contours in **A** and **D**, and by colored horizontal line segments in **B, C, E** and **F**.

Figure 4B depicts lateralized power in the alpha band for Same- and Opposite trials separately. These data are largely in line with the observations regarding the grand-average data: Both conditions showed strong, contralateral alpha suppression in response to the presentation of the onset marker, though this qualitatively seemed to be more pronounced for Same-trials. Both conditions also seemed to show anticipatory shifts in alpha suppression reflecting the location of the upcoming offset marker: Same-trials showed significant contralateral suppression from −709ms to 766ms, and Opposite trials seemed to shift polarity leading up to the offset marker, resulting in a significant negative cluster from −55 to 639ms. There were, however, no significant differences between data from these two conditions.

Figure 4C depicts the same time courses of lateralized alpha power separately for correct and incorrect responses. As with the ERPs, no significant differences were found between the two conditions in either of the epochs, and both trial types had very similar signatures of lateralized alpha suppression. Of note, the lateralized alpha suppression was not marked by any significant cluster on incorrect trials, even though it was qualitatively very similar to that on correct trials.

Figures 4D–F depict the same analyses for data around the onset and offset markers of Interval 2. Across these plots, neither of the conditions gave any indication of lateralized suppression related to the presentation of these markers. In the grand-average time-frequency spectra (D), no significant clusters were found. Same- and Opposite trials (E) showed significant, opposite lateralization responses during the memory delay, with a siginificant difference between them early in this interval, but these lateralized responses seemed to resolve right before the onset of Interval 2. No other significant differences were found between these two conditions. Similarly, there were no signs of lateralized alpha suppression for data from correct- or incorrect trials, and no differences between these two conditions.

To conclude, the time-frequency analyses generally aligned with the ERP analyses: we found lateralized alpha suppression indicative of attentional orienting in response to the onset and offset markers of Interval 1. Of note, lateralized responses already arose in anticipation of the predictable offset marker, suggestive of early attentional shifts to the anticipated marker location. However, surrounding the markers of Interval 2, there was no indication that these lateral locations were attended or retrieved again.

### Classification

The classification analyses explored whether during comparison, marker locations were maintained, retrieved or reactivated in a manner that was not captured by analyses above, which focused on lateralization in occipito-parietal sites. To this end, we trained classifiers to decode the location of onset or offset cues of Interval 1, based on the broadband signal from all 64 channels. The results of this analysis are largely in line with our findings from ERP- and time-frequency analyses.

Figure 5 depicts, for each epoch, the performance of classifiers trained on the overall dataset, as well classifiers trained and tested separately on ‘Same’ or ‘Opposite’ trials. Time courses of classification performance depicted that both the onset- and offset marker of Interval 1 could be decoded from the data epochs around their presentation, both for a sustained period of time with an AUC significantly above chance from 109 to 984ms and from 109 to 1031 ms for onset and offset epochs, respectively. Separately classifying ‘Same’ and ‘Opposite’ trials yielded virtually identical patterns of results.

**Figure 5.**
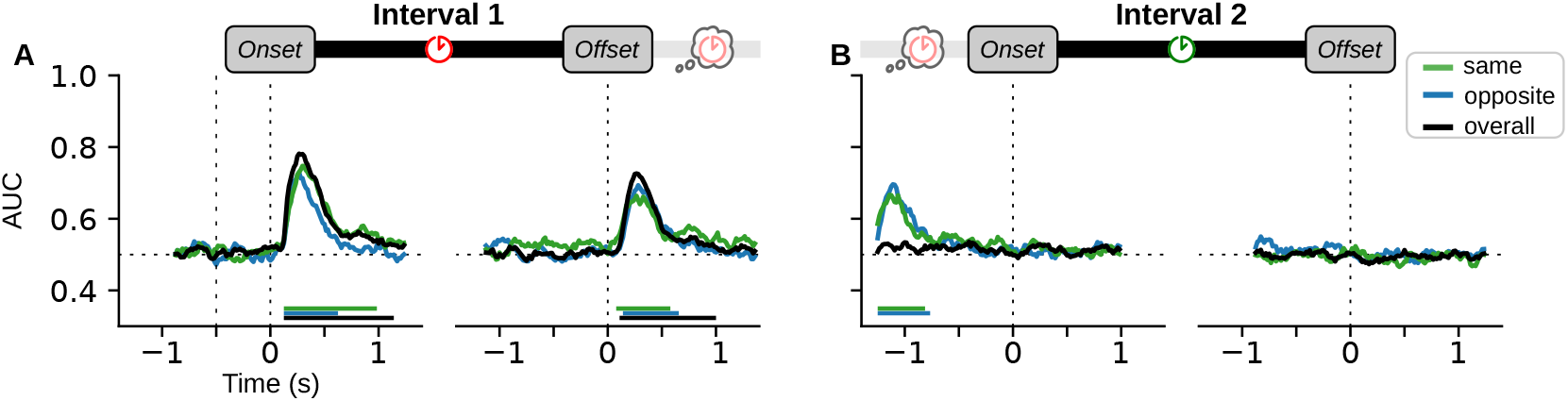
Classifier performance (Area Under the Curve) derived from 10-fold cross validation on either only ‘Same’ blocks, only ‘Opposite’ blocks, or all trials together (overall). **A** Classifiers trained to decode the Interval 1 onset marker location (left) or the offset marker location, on corresponding data from Interval 1 **B** Classifiers trained and tested on data from Interval 2. For these epochs, the location of the onset- or offset marker of Interval 1 were the classification target.

As can be seen from the overall classification scores during the memory delay, multivariate EEG robustly and persistently reflects the location of the offset marker, which gradually dissipates as the onset marker draws closer (Figure 5A, right plot). However, when the same data are used to decode the onset marker location, leading up to the onset of Interval 2, classification scores suggest that this location is not reflected in multivariate data (Figure 5B, left). It may seem surprising that we find significant decoding above chance in this latter analysis when Same- and Opposite trials are considered separately, but not when both trials are pooled together. However, recall that in each of these conditions, the Interval 1 offset marker location (and its associated visual response) are fully predictive of the onset marker location. Therefore, these increments in classifier performance essentially reflect the visual response to the offset marker.

After the onset of the centrally presented Interval 2, none of the classifiers were able to accurately determine the marker locations of Interval 1. No other significant clusters of decoding performance were found.

To conclude, the classification results were in line with the ERP and time-frequency analyses: Results again indicated that marker locations were persistently represented after they had been presented as markers for Interval 1, but that these locations were not memorized, retrieved, or otherwise represented during comparison.

## Discussion

Many studies exploring the dynamics of working memory representations have exploited the finding that visual features in working memory appear to be bound to the location at which they were presented (de Vries et al., 2017; Kuo et al., 2011; Poch et al., 2014; van Ede et al., 2019). That is, maintaining or retrieving non-spatial visual information has been found to evoke clear neural markers in the ERP and time-frequency spectra that are reflective of the location of presentation. Here, we investigated whether temporal information is similarly bound to spatial information during encoding, maintenance and retrieval, in order to explore the dynamics of temporal memory. Participants perceived an interval presented by means of lateralized start- and end-marker stimuli, and we found that the encoding of the interval indeed evoked spatial neural signatures typically associated with working memory retention. However, when this interval was subsequently compared to an interval with central markers, no signs of lateralization were found; neither in the ERP, nor in the time-frequency spectrum, nor could spatial information otherwise be decoded using multivariate pattern analysis. These results suggest that while memory for visual features such as location might play a role during initially timing an interval, spatial information is not involved when the interval is subsequently used for comparison.

The start marker of the first interval produced lateralized ERP components, lateralized alpha suppression, and sustained robust classification scores that are highly comparable to those found in typical studies on visual working memory encoding (Carlisle et al., 2011; de Vries et al., 2017; Gunseli et al., 2014; Van Driel et al., 2017; Wolff et al., 2017), which we interpret as signatures of working memory involvement in this timing task. One notable difference compared to those studies is that from 300 to 400ms, the results reveal a contralateral positivity where we had anticipated to find an N2pc-component. One could suspect that this discrepancy stems from the laterally unbalanced displays used in our design: in most working memory studies the lateralized memorandum is presented alongside a non-target in the opposite hemifield. Indeed, unilaterally presented, highly salient stimuli have been shown to evoke a so-called P2pc in this time window (Casiraghi et al., 2013). However, the present results are very similar to those in early investigations of the N2pc, in which displays could be similarly ‘unbalanced’ (Hickey et al., 2009). Based on these similarities, it seems that this positivity could also reflect a component known as a distractor positivity (*P*_*D*_, see also Burra & Kerzel, 2014; Sawaki & Luck, 2010, 2013). This component could reflect how participants actively suppressed this salient onset in order to maintain central fixation. Critically, however, despite the initial ‘distractor status’ of the marker, this positive component still reversed into a sustained negativity that reflected a CDA.

The neural responses elicited by the end marker of the first interval were largely similar to those caused by its onset. Notably, the end marker resulted in a CDA-component that lasted well into the delay period, qualitatively even seeming to persist into the second interval. If the CDA is caused by the working memory processes involved in timing, this raises the question why another CDA would arise after the ‘timing work’ is already done. One possible explanation would be that this signature reflects maintenance of the relevant duration during the delay interval. If this were the case, however, we would have anticipated a CDA-difference between ‘Same’ and ‘Opposite’ trials, where the former trial type would be much stronger associated with its location than the latter. Another explanation would be that the delay period in itself is actually an interval that is being timed, so as to optimally anticipate the start of the comparison interval.

Despite these lateralized neural signatures found during Interval 1, our results indicate that the central markers of Interval 2 did not convey any information regarding the preceding locations of Interval 1. One theoretical account that fits these results poses that while time perception may result from distributed intrinsic circuits, these circuits might subsequently project to more centralized ‘readout neurons’ that allow for the comparison of intervals across distributed clocks (Bakhurin et al., 2017; Laje & Buonomano, 2013; Paton & Buonomano, 2018). Under this model, a more generic, stimulus-independent representation of duration would be extracted at the offset of Interval 1, which is subsequently maintained in memory and used for comparison. As such, the spatial code might be involved in the initial perception of time, but not retained or retrieved when the interval is compared later on.

Another, subtly different account would pose that perception- and comparison of duration make use of different representations of the Interval 1 duration, and that transforming one into the other is an active process. Such transformations of have been proposed to be used for visual search (Myers et al., 2017), where multiple items can be perceived and stored in working memory, after which only one relevant item is actively transformed into a ‘template’ that guides attentional selection without interference from others (see also Chatham et al., 2014; Olivers et al., 2011; Ort & Olivers, 2020). However, one key difference is that for visual search there is strong evidence that transformed representations still make use of the spatial code (de Vries et al., 2018; de Vries et al., 2017, 2019), whereas for temporal information this association appears to be absent. If temporal comparison uses such a transformation, it may be the case that this only takes place after the memory delay, which might account for the persistent lateralization observed throughout the memory delay. However, it seems likely that such a transformation would take place before the end of Interval 2, given the ample evidence that temporal discrimination judgments can be made before the offset of the comparison interval (Balcı & Simen, 2014; Bueno & Cravo, 2020; Macar & Vidal, 2003).

As such, our results do not unequivocally support either dedicated or intrinsic clock models. On the one hand, the dynamics observed during Interval 1 are in line with what one would predict following intrinsic models of time perception. However, the observation that the spatial code appears to be abandoned during comparison largely falls in line with dedicated models. Of note, our behavioral data yielded unexpected support for intrinsic clock models: the shift in perceived time between ‘Same’ and ‘Opposite’ blocks indicates that a relatively subtle difference in the manner of presentation can have a profound impact on the temporal percept (in line with Droit-Volet & Meck, 2007; Eagleman & Pariyadath, 2009; Johnston et al., 2006; Matthews, 2011). These results might align with how the perception of time and space are closely intertwined, as has been argued before (Bueti & Walsh, 2009; Burr et al., 2010; Dehaene & Brannon, 2010). This behavioral bias coincided with ERP differences in response to the offset markers of ‘Same’ and ‘Opposite’ markers, a finding that warrants more thorough exploration beyond the scope of the present study. We discuss this observation in more detail in a companion article (Kruijne et al., 2020).

The mechanisms by which we perceive sensory events, and the experience of the timing of these events are intertwined by necessity. The present results offer new, important insights into the interplay of stimulus representations and temporal representations during perception, maintenance, and subsequent comparisons of interval durations. Despite converging evidence that many features of visually presented stimuli are bound to spatial information when encoded in working memory, we found no evidence that maintaining and retrieving temporal information relied on such an associative link. These results offer critical constraints for models that not only aim to measure time, but also to subsequently use that measurement in upcoming goal-directed behaviors. In other words, these findings offer important considerations for theories of temporal cognition beyond the perception of time.

## Acknowledgements

The authors would like to thank Philippa Johnson for her help in data acquisition, and thank Joram van Driel for his help with analyses and experimental design.

## Footnotes

1 Another approach would be to subtract pre-stimulus baseline activity for each marker separately. That approach is better suited to isolate effects in the transient evoked responses post-stimulus, but is likely to obscure potential slow-wave lateralization differences that might still be present right before stimulus presentation (like the CDA). We have ran analyses where epochs were re-baselined as such, but these did not lead to different conclusions

2 in this and all subsequent descriptions, we will treat adjacent clusters as one ‘effect’ wherever they are less than 50ms apart and denote effects in the same direction.

